# Integrating water, sanitation, handwashing, and nutrition interventions to reduce child soil-transmitted helminth and *Giardia* infections: a cluster-randomized controlled trial in rural Kenya

**DOI:** 10.1101/464917

**Authors:** Amy J. Pickering, Sammy M. Njenga, Lauren Steinbaum, Jenna Swarthout, Audrie Lin, Benjamin F. Arnold, Christine P. Stewart, Holly N. Dentz, MaryAnne Mureithi, Benard Chieng, Marlene Wolfe, Ryan Mahoney, Jimmy Kihara, Kendra Byrd, Gouthami Rao, Theodora Meerkerk, Priscah Cheruiyot, Marina Papaiakovou, Nils Pilotte, Steven A. Williams, John M. Colford, Clair Null

## Abstract

**Background.:** Helminth and protozoan infections affect >1 billion children globally. Improved water, sanitation, handwashing, and nutrition could be more sustainable control strategies for parasite infections than mass drug administration (MDA), while providing other quality of life benefits.

**Methods and Findings.:** We enrolled geographic clusters of pregnant women into a cluster-randomized controlled trial that tested six interventions: disinfecting drinking water(W), improved sanitation(S), handwashing with soap(H), combined WSH, improved nutrition(N), and combined WSHN. We assessed intervention effects on parasite infections by measuring *Ascaris lumbricoides*, *Trichuris trichiura*, hookworm, and *Giardia duodenalis* among individual children born to enrolled mothers and their older siblings (ClinicalTrials.gov NCT01704105). We collected stool specimens from 9077 total children in 622 clusters, including 2346 children in control, 1117 in water, 1160 in sanitation, 1141 in handwashing, 1064 in WSH, 1072 in nutrition, and 1177 in WSHN. In the control group, 23% of children were infected with *Ascaris lumbricoides*, 1% with *Trichuris trichuria*, 2% with hookworm and 39% with *Giardia duodenalis*. After two years of intervention exposure, *Ascaris* infection prevalence was 18% lower in the water treatment arm (95% confidence interval (CI) 0%, 33%), 22% lower in the WSH arm (CI 4%, 37%), and 22% lower in the WSHN arm (CI 4%, 36%) compared to control. Individual sanitation, handwashing, and nutrition did not significantly reduce *Ascaris* infection on their own, and integrating nutrition with WSH did not provide additional benefit. *Trichuris* and hookworm were rarely detected, resulting in imprecise effect estimates. No intervention reduced *Giardia*. Reanalysis of stool samples by quantitative polymerase chain reaction (qPCR) confirmed the reductions in *Ascaris* infections measured by microscopy in the WSH and WSHN groups. Lab technicians and data analysts were blinded to treatment assignment, but participants and sample collectors were not blinded. The trial was funded by the Bill & Melinda Gates Foundation and USAID.

**Conclusions.:** Our results suggest integration of improved water quality, sanitation, and handwashing could contribute to sustainable control strategies for *Ascaris* infections, particularly in similar settings with recent or ongoing deworming programs. Water treatment alone was similarly effective to integrated WSH, providing new evidence that drinking water should be given increased attention as a transmission pathway for *Ascaris*.

## Introduction

Intestinal soil-transmitted helminth (STH) infections, including *Ascaris lumbricoides*, *Trichuris trichiura*, and hookworm, and the protozoa *Giardia duodenalis* are common parasitic infections among children in low-resource settings and neglected tropical diseases. Globally, STH are estimated to affect 1.45 billion people(1), while *Giardia* has been cited as the most common enteropathogen in low-income countries(2). STH and *Giardia* infections can result in poor absorption of nutrients and weight loss(3,4). There is some evidence that STH and *Giardia* infections, even when asymptomatic, may contribute to growth faltering and impaired cognitive development(5-8). Longitudinal cohort studies in Bangladesh and Brazil have identified early infection with *Giardia* as a risk factor for stunting among children(7,9). In Peru, children with multiple *Giardia* infections per year during the first two years of life had lower cognitive function scores at age 9 than children with one or fewer *Giardia* infections(10). The effect of child STH infections on child growth, cognitive development, and school performance has been mixed and strongly debated by experts, with some suggesting additional evidence is needed(5,6,11).

School-based mass drug administration (MDA) campaigns have been the cornerstone of the global strategy to control STH infections; however, high reinfection rates limit the ability of MDA to achieve sustained reduction in STH infection prevalence(12). *Ascaris*, *Trichuris*, *Giardia*, and *Ancylostoma duodenale* are primarily transmitted through the fecal-oral ingestion route, although *Ancylostoma duodenale* as well as *Necator americanus* can be transmitted transdermally. A meta-analysis of studies from settings with medium-to-high endemic STH prevalence identified an average reinfection rate for *Ascaris* at 12 months at 94% of baseline prevalence, while the average 12-month reinfection rates for *Trichuris* and hookworm were 82% and 57%, respectively(13). To achieve elimination of STH transmission, it has been suggested that MDA control efforts may need to be integrated with improved water, sanitation, and handwashing(14). Control of *Giardiasis* has historically relied on drug treatment after diagnosis as well as exposure prevention by water treatment and improved sanitation, but zoonotic transmission can complicate exposure prevention(15). Recent systematic reviews suggest that improved water, sanitation, and handwashing can reduce the odds of STH and *Giardia* infections, though the quality of the evidence base remains poor and consists almost exclusively of observational analyses(16,17).

An individual’s susceptibility to STH and *Giardia* infection is influenced by exposure and immune response. A recent systematic review concluded that there was some evidence that nutritional supplementation decreases the risk of infection or reinfection with STH, but studies have been of low quality(18). Plausible mechanisms by which nutrition might reduce STH or *Giardia* infection are through improvements in effective immune response including repair of cell damage caused by parasite infection, and through changes to the gut microbiome(19,20).

We conducted a cluster-randomized controlled trial in rural Kenya to assess the effects of water, sanitation, handwashing, and nutrition interventions delivered alone and in combination on child parasite infections. STH and *Giardia* infections were pre-specified as trial outcomes before the trial began(21). In a separate paper, we reported the effects of the interventions on child growth and diarrhea(22). The trial’s nutrition intervention was the only component that improved child growth, but none of the interventions reduced diarrhea(22). Here, we report intervention effects on *Ascaris*, *Trichuris*, hookworm, and *Giardia* infections measured after two years of intervention exposure.

## Materials and Methods

### Study design

The trial protocol and detailed methods are published(21). The trial was registered at ClinicalTrials.gov, identification number: NCT01704105. The study protocol was approved by the Committee for the Protection of Human Subjects at the University of California, Berkeley (protocol number 2011-09-3654), the Institutional Review Board at Stanford University (IRB-23310), and the Scientific and Ethics Review Unit at the Kenya Medical Research Institute (protocol number SSC-2271). Innovations for Poverty Action (IPA) enrolled participants, implemented the intervention delivery, and collected the data. Mothers provided written informed consent for themselves and their children.

Clusters of eligible pregnant women each were randomized by geographic proximal blocks into one of eight study arms: chlorine treatment of drinking water (W); improved sanitation including provision of toilets with plastic slabs and hardware to manage child feces (S); handwashing with soap (H); combined WSH; infant and young child feeding counseling plus small-quantity lipid-based nutrient supplements (N); combined WSHN; a double-sized active control, and a passive control arm. Children in the passive control arm were purposively excluded from parasitology measurement (Figure 1).

**Figure 1.**
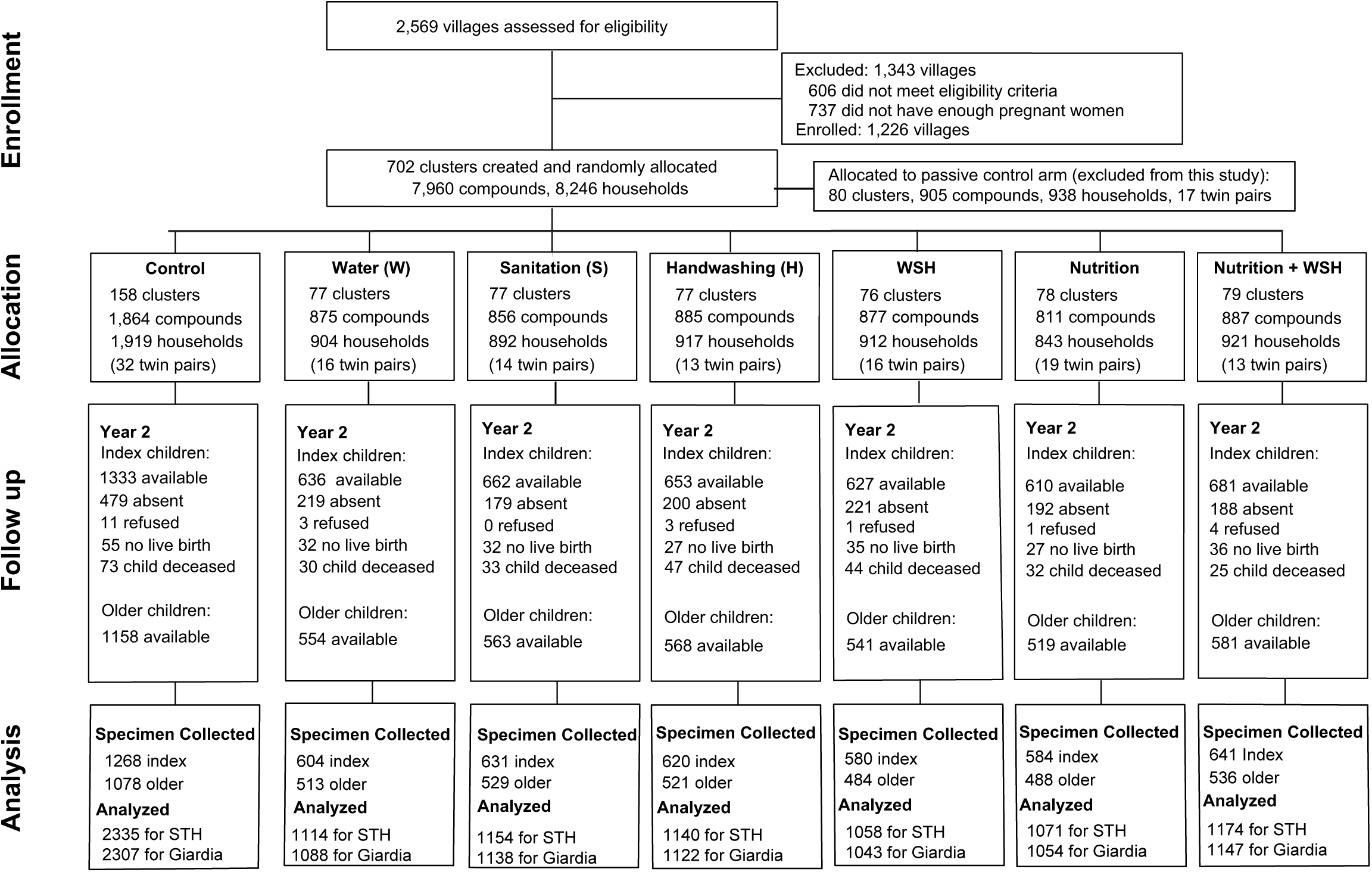
Trial profile.

We conducted a cluster-randomized trial because components of the intervention promotion activities were at the community level and there could have been behavior and infectious disease interactions between neighboring households. Villages were eligible for selection into the study if they were rural, the majority of the population lacked access to piped water supplies, and there were no other ongoing WSH or nutrition programs. Within selected villages, a census was conducted to identify eligible pregnant women in their second or third trimester that planned to continue to live at their current residence for the next year. Since interventions were designed to reduce child exposure to pathogens through a cleaner environment and exclusive breastfeeding, we enrolled pregnant women to allow time for intervention delivery to occur prior to or as close to birth as possible. Clusters were formed from 1-3 neighboring villages and had a minimum of six pregnant women per cluster after the enrollment survey. Children born to enrolled pregnant mothers were considered “index” children. Outcomes were assessed after two years of intervention exposure among index children, including twins, as well as among one older child in the index child’s compound to understand the effect of the interventions on both preschool aged and school aged children. The older child was selected by enrolling the youngest available child within the age range of 3-15 years old, with priority for a sibling in the index child’s household.

### Baseline survey

A survey at enrollment measured household socioeconomic characteristics and demographics (maternal age, maternal education, electricity access, type of floor, number of people in the household), as well as water, sanitation, and handwashing infrastructure and behaviors (type of water source, reported water treatment, defecation location, type of toilet, presence of water and soap at a handwashing station). In addition, at study enrollment we measured *Giardia*, *Entamoeba histolytica* and *Cryptosporidium spp*. among children residing in study compounds between 18 and 27 months of age (the projected age range for index children at the end of the study) to assess baseline prevalence of these pathogens. STH were not measured at enrollment among these proxy children because it was not logistically feasible to deworm infected children at baseline. We also collected 100ml samples from primary drinking water sources accessed by study households and household stored drinking water (if available). We transported the samples on ice to field labs and enumerated *Escherichia coli* in each sample by membrane filtration followed by culture on MI media.

### Randomization and blinding

A few weeks after enrollment, clusters were randomly assigned to intervention arms at the University of California, Berkeley by an investigator independent of the field research team (BFA) using a random number generator. Groups of nine, geographically adjacent clusters were block-randomized into the six intervention arms, the double-sized active control arm, and the passive control arm (the passive control arm was not included in the parasite assessment). Participants and other community members were informed of their intervention group assignment after the baseline survey. Blinding (masking) participants was not possible given the nature of the interventions. Data and stool sample collectors were not informed of the cluster intervention assignment, but could have inferred treatment status by observing intervention hardware. Lab technicians were blinded to intervention status. Two authors (AJP and JS) independently replicated the statistical analyses while blinded to intervention status.

### Intervention delivery

Intervention delivery began <3 months after enrollment. In the water intervention arms (W, WSH, WSHN), community health promoters encouraged drinking water treatment with chlorine (liquid sodium hypochlorite) using either manual dispensers installed at the point-of-collection (community water source) in study villages or using bottled chlorine provided directly to households every 6 months. In the sanitation arms (S, WSH, WSHN), households received new latrines or existing latrines were upgraded and improved by installing a plastic slab that included a lid. All households in sanitation arm study compounds were provided with a child potty for each child <3 years as well as a “sani-scoop” to remove animal and human feces from the compound. In the handwashing arms (H, WSH, WSHN), households were provided with two handwashing stations—near the latrine and the cooking area. Stations included dual foot-pedal operated jerry cans that could be tipped to dispense either soapy water or rinse water. Households were responsible for keeping the stations stocked with rinse water, and community health promoters refilled soap regularly. In the nutrition arms (N, WSHN), small quantity lipid-based nutrient supplements (LNS) were provided to children from 6-24 months of age. Children received monthly rations of LNS for addition to complementary foods twice per day. Nutrition messaging included promoting dietary diversity during pregnancy and lactation, early initiation of breastfeeding, exclusive breastfeeding from 0-6 months, continued breastfeeding through 24 months, timely introduction of complementary foods, dietary diversity for child feeding, and child feeding during illness.

Community health promoters were nominated by mothers in the community and trained to provide intervention-specific behavior change activities and instructions on hardware use or provision of nutrition supplements. They were also trained to measure child mid-upper arm circumference to identify and provide referrals for potential cases of severe acute malnutrition. Each intervention consisted of a comprehensive behavior change package of key messages; visual aids in the form of flip charts, posters, and reminder cue cards; interactive activities with songs, games, or pledges to commit to practice target behaviors; and the distribution of arm-specific hardware, products, or supplements. Households in the active control group received visits from promoters to measure child mid-upper arm circumference and provide malnutrition referrals, but did not receive any intervention related hardware or messaging. Promoters were instructed to visit households monthly. Key messages and promoter materials are available at https://osf.io/fs23x/.

Adherence to the interventions was measured during unannounced household visits after one year and two years of intervention exposure (see SI).

### Measurement of parasite infections

We measured parasite infections approximately 27 months post-enrollment (which equates to a minimum of 24 months of intervention exposure since intervention hardware was delivered <3 months of enrollment). Stool samples were collected from index children and older children in sterile containers and transported on ice to the closer of two central field labs located in Kakamega or Bungoma. Field staff revisited households up to 3 times to collect stool samples. *Ascaris lumbricoides*, *Trichuris trichiura*, and hookworm eggs were immediately enumerated (same day) by double-slide Kato Katz microscopy with 41.7 mg templates. Both slides created from each stool sample were counted by a trained parasitologist, and two different parasitologists counted each slide from the same sample. A supervisor with expertise in STH egg identification reviewed 10% of all slides and any discrepancies were corrected. STH egg counts were averaged for analysis if both slides from one stool sample were positive. Two aliquots of stool (one mixed with ethanol) were transported on dry ice to the Eastern and Southern Africa Centre of International Parasite Control laboratory at KEMRI in Nairobi, Kenya for further analysis.

One aliquot was analyzed by monoclonal enzyme-linked immunosorbent assay (ELISA) assay (*Giardia II^TM^*, Alere International Limited, Galway, Ireland) for the presence or absence of *Giardia duodenalis* cysts. Samples were measured by ELISA in duplicate; if there was a discrepancy between duplicates, the sample was re-run. DNA was extracted from the other aliquot (preserved in ethanol) for stool samples collected from children in the control, WSH, and WSHN groups. Four qPCR assays were run in duplicate on each sample to detect the following targets: *Necator americanus*, *Ancylostoma duodenale*, *Trichuris trichiura*, and *Ascaris lumbricoides* (see SI for further details)(23).

### Outcomes

STH and *Giardia* infections were pre-specified outcomes in the parent WASH Benefits trial prior to the start of data collection; see Figure 3 in Arnold and others (21). Parasite infections were measured two years after the start of intervention activities. The main indicators of parasite infections were prevalence of each individual STH infection, any STH infection, and the prevalence of *Giardia* infection among index and older children from the same compound. Additional indicators of parasite infections included intensity of *Ascaris*, *Trichuris*, and hookworm measured in eggs per gram (epg) of feces; intensity binary category of *Ascaris* infection measured as low intensity (1-5000 epg) or moderate/high intensity (>5000 epg) infection following World Health Organization (WHO) cutoffs; prevalence of co-infection with two or three STH; and prevalence of co-infection with *Giardia* and any STH. The trial’s original protocol included *Entamoeba histolytica* and *Cryptosporidium spp*. as additional protozoan endpoints. At enrollment, *Giardia* prevalence was 40% among 535 children 18-27 months old in study compounds, while *Cryptosporidium Spp.* prevalence was 1% and *E. histolytica* prevalence was 0%. We determined the extremely low prevalence made these trial endpoints futile due to limited statistical power, and since each required a separate assay on the ELISA platform, the study’s steering committee decided to not test for them at follow-up.

### Sample size calculations

All households in all clusters enrolled into the main trial were invited to participate in the measurement of parasite infections. The main trial was powered for a minimum detectable effect of 0.15 in length-for-age *Z* score and a relative risk of diarrhea of 0.7 or smaller for a comparison of any intervention with the double-sized control group, assuming a type I error (α) of 0·05 and power (1–β) of 0.8, a one-sided test for a two-sample comparison of means, and 10% loss to follow-up. This led to a planned design of 100 clusters per arm and 10 index children per cluster. Given this design and a single, post-intervention measure, we estimated that the trial’s sample size would be sufficient at 80% power with a two-sided ? of 0.05 to detect a relative reduction of 18% in infection prevalence of any parasite. Our minimum detectable effect calculations assumed 50% prevalence in the control arm, a village intraclass correlation (ICC) of 0.14, two children measured per enrolled household (index child plus an older sibling), and 70% successful stool collection and analysis.

### Statistical analysis

All statistical analyses and comparisons between arms (W, S, H, WSH, N, WSHN compared to active-control) were pre-specified prior to unblinding of investigators and published with a time-stamp on the Open Science Framework (OSF) (https://osf.io/372xf/). Replication scripts and data are also provided at the same link. Our alternative hypothesis for all comparisons was that group means were not equal (two-sided tests). We estimated unadjusted and adjusted intention-to-treat effects between study arms using targeted maximum likelihood estimation (TMLE) with influence curve-based standard errors that treated clusters as independent units and allowed for outcome correlation within clusters(24,25). Our parameters of interest for dichotomous outcomes were prevalence ratios. Our parameter of interest for helminth intensity was the relative fecal egg count reduction. We calculated the relative reduction using both geometric and arithmetic means. We did not perform statistical adjustments for multiple outcomes to preserve interpretation of effects and because many of our outcomes were correlated(26). We estimated adjusted parameters by including variables that were associated with the outcome to potentially improve the precision of our estimates. We pre-screened covariates (see SI for full list) to assess whether they are associated (P-value <0.2) with each outcome prior to including them in adjusted statistical models. We conducted subgroup analyses to explore effect modification on *Ascaris* and *Giardia* infection presence for the following factors: index child status, consumed deworming medicine in past 6 months (*Ascari*s only), consumed soil in past week (index children only), >8 people in compound, and if defecation occurred on the same day as stool collection. Statistical analyses were conducted using R version 3.3.2 (www.r-project.org).

## Results

### Enrollment

Pregnant women were enrolled into the cluster-randomised controlled trial from Kakamega, Bungoma, and Vihiga counties in Kenya’s western region. Enrollment occurred between November 2012 - May 2014; 8246 pregnant women were enrolled. Clusters with an average of 12 eligible pregnant women each were randomized by geographic proximal blocks into one of eight study arms: chlorine treatment of drinking water (W); improved sanitation including provision of toilets with plastic slabs and hardware to manage child feces (S); handwashing with soap (H); combined WSH; infant and young child feeding counseling plus small-quantity lipid-based nutrient supplements (N); combined WSHN; a double-sized active control, and a passive control arm. Children in the passive control arm were purposively excluded from parasitology measurement (Figure 1). Parasite infections were measured among children born to enrolled pregnant mothers (index children) as well as their older siblings.

Enrollment characteristics of the study population were similar between arms (Table S1). Most households accessed springs or wells as their primary drinking water source. In the control group, 24% of households accessed unprotected water sources, such as springs, dug wells, and surface water. The microbial quality of drinking water was very poor, as has been reported previously for this study area(27); 96% (n=1829) of source water samples and 94% (n=5959) of stored drinking water samples contained *Escherichia coli* contamination. Most (82%) households owned a latrine, but only 15% had access to a latrine with a slab or ventilation pipe (Table S1). Soap and water availability for handwashing at a designated handwashing location was low (<10%).

### Indicators of intervention uptake

After one year of intervention, 89-90% of households that received the sanitation intervention had access to an improved latrine (compared to 18% in active-control arm) and 79-82% of these had access to an improved latrine after two years of intervention. In the water intervention arms, 40-44% of households had a detectable chlorine residual in their stored drinking water at the one-year follow up (compared to 3% of control households) and 19-23% had chlorine detected after two years. 76-78% of households that received the handwashing intervention had soap and water available at a handwashing station (compared to 12% in the control arm) after one year and this decreased to 19-23% at year two. Consumption of LNS sachets by children in the nutrition arms was 95-96% of the expected two sachets per day at the one-year follow up and 114-116% of expected at the 2-year follow up (>100% is possible because additional LNS packets were delivered in case of future delivery delays) (Tables S2 & S3).

### Infection prevalence

Soil-transmitted helminth and *Giardia* infections were measured after two years of exposure to the interventions. We collected stool specimens from 9077 children aged 2-15 years old at the two-year survey during January 2015 – July 2016; including 4928 index children (median age in years: 2.0, interquartile range (IQR): 1.9, 2.1) and 4149 older children (median age in years: 5.0, IQR: 4.2, 6.4) residing in an index child’s compound (Figure 1). A total of 2346 children in 158 control clusters, 1117 children in 77 water clusters, 1160 children in 77 sanitation clusters, 1141 children in 77 handwashing clusters, 1064 children in 76 WSH clusters, 1072 children in 78 nutrition clusters, and 1177 children in 79 WSHN clusters provided stool specimens. Stool specimens were successfully collected from 95% (4928 of 5202) of available index children and from 93% (4149 of 4484) of available older children two years after intervention delivery (Figure 1 shows number of children not available due to no live birth, death, refusal, or absent; Table S7 shows characteristics of children lost to follow up). In the control group 22.6% of children were infected with *Ascaris* (ICC: 0.10), 2.2% with hookworm (ICC: 0.04), 1.2% with *Trichuris* (ICC: 0.07) (measured by Kato-Katz microscopy), and 39% with *Giardia* (measured by enzyme-linked immunosorbent assay)(Table S4). *Ascaris* infection prevalence was similar for index children (22.8%) and older children (22.3%) in the control group (Table S6). Caregivers reported that 39% of index children and 10% of older children had consumed soil in the past 7 days.

### Effect of interventions on parasite infection prevalence

Infection prevalence of each STH, any STH, and *Giardia* was compared between each intervention group (W, S, H, WSH, WSHN) and the double-sized active control group (C); see methods for further details of the analysis. Compared to the control group, *Ascaris* infection prevalence was 18% lower in the water arm (Prevalence Ratio [PR]: 0.82, 95% Confidence Interval [CI] 0.67, 1.00), 22% lower in the combined WSH arm (PR: 0.78, 95% CI 0.63, 0.96), and 22% lower in the WSHN arm (PR: 0.78, 95% CI 0.64, 0.96) (Figure 2, Table S4). Sanitation, handwashing, and nutrition did not significantly reduce *Ascaris* infection on their own (Figure 2). The combined WSH intervention reduced infection with any STH by 23% (PR: 0.77, 95% CI 0.63, 0.95) and the combined WSHN intervention reduced infection with any STH by 19% (PR: 0.81, 95% CI 0.66, 0.98) (Table S4). No interventions significantly reduced the prevalence of hookworm and *Trichuris*, though the low prevalence in the control arm meant that any reduction due to intervention would be difficult to detect in the trial (Table S4). No interventions reduced *Giardia* prevalence (Figure 2).

**Figure 2.**
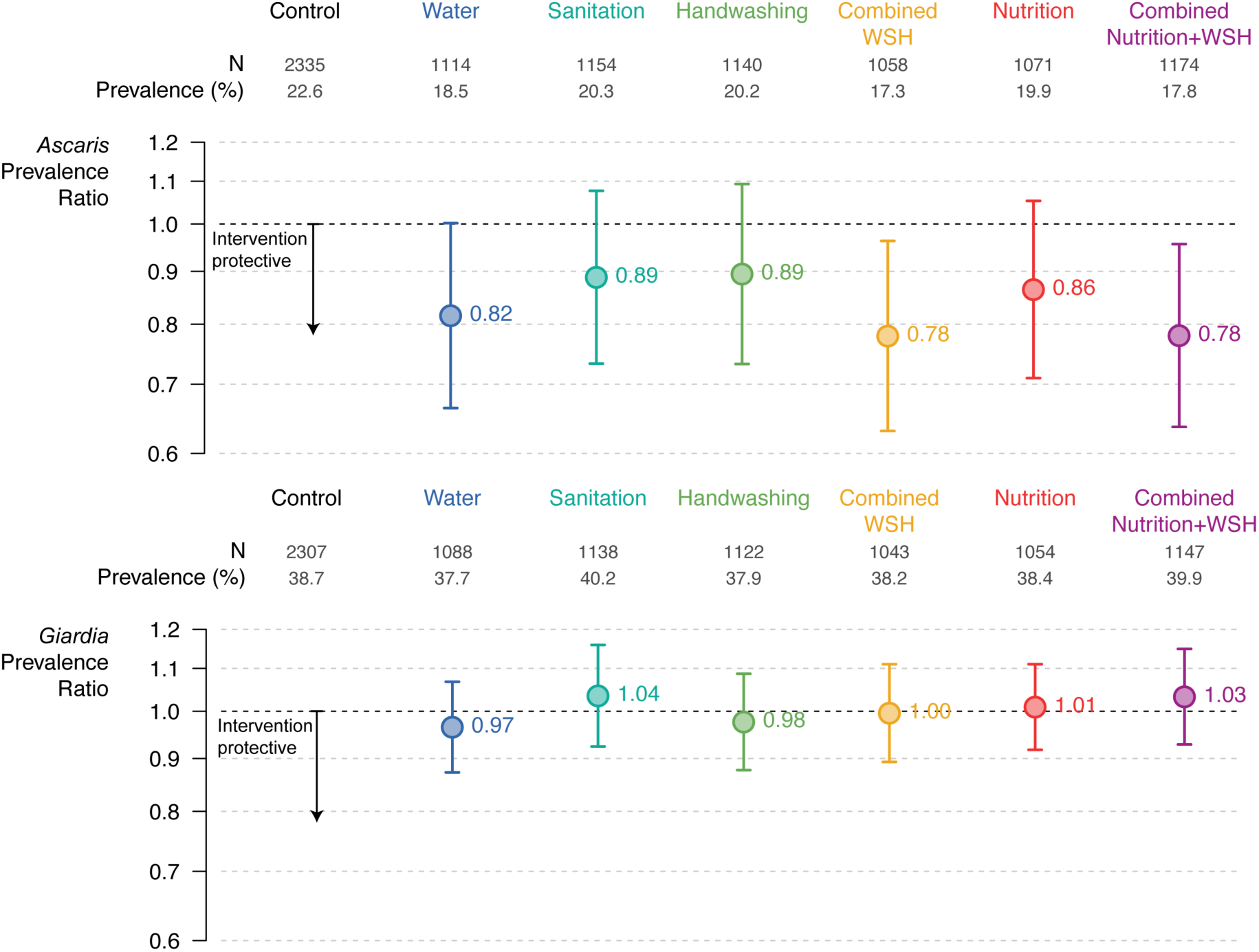
Effect of the interventions on infection with *Ascaris* and *Giardia.* Prevalence ratios estimated by targeted maximum likelihood estimation. Error bars show 95% confidence intervals for the prevalence ratios.

We re-analyzed all stool samples collected from children enrolled in the control, combined WSH, and WSHN arms by quantitative polymerase chain reaction (qPCR) to validate our estimates based on microscopy measurements. These three arms were selected for the qPCR subset analysis prior to unblinding of investigators to results and were chosen based on the hypothesis that these arms would be the most likely to have low-intensity STH infections if any of the interventions were effective. qPCR analyses resulted in almost identical intervention effect estimates to those based on microscopy (Figure 3, Table S8). Compared to the control group, *Ascaris* infection prevalence was 21% lower (PR: 0.79, 95% CI 0.64, 0.97) in the WSN group and 23% lower (PR: 0.77, 95%CI 0.64, 0.93) in the WSHN group. We also did not detect any significant effects of the interventions on *Trichuris* or hookworm infections using qPCR data (Table S8).

**Figure 3.**
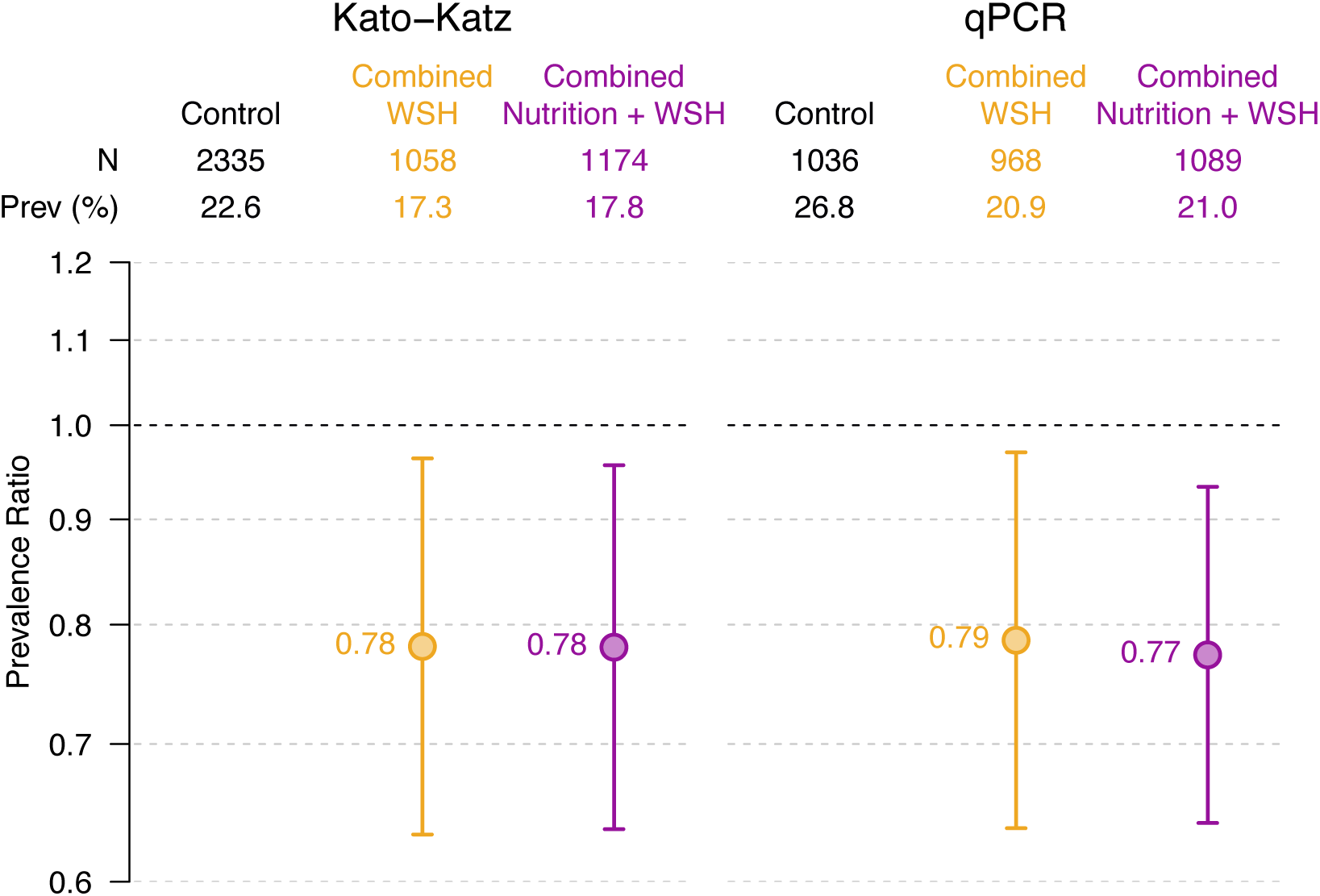
Effect of the combined interventions on infection with *Ascaris* estimated with Kato-Katz microscopy (left) and by qPCR (right). Prevalence ratios estimated by targeted maximum likelihood estimation. Error bars show 95% confidence intervals for the prevalence ratios.

### Effect of interventions on infection intensity

*Ascaris* infection intensity was lower in children in the water arm (fecal egg count reduction with geometric means [FECR]: −16%, 95% CI −32%, −1%), the WSH arm (FECR: −19%, 95% CI −33%, −5%), and the WSHN arm (FECR: −18%, 95% CI −32%, −4%) compared to the control arm; FECR with arithmetic means showed similar results (Table 1). The prevalence of heavy/moderate intensity *Ascaris* infections was 10.0% in the water arm, 10.9% in WSH, and 10.3% in WSHN compared to 12.7% in the control arm; these differences were not statistically significant at the 95% confidence level (Table S4).

**Table 1.**
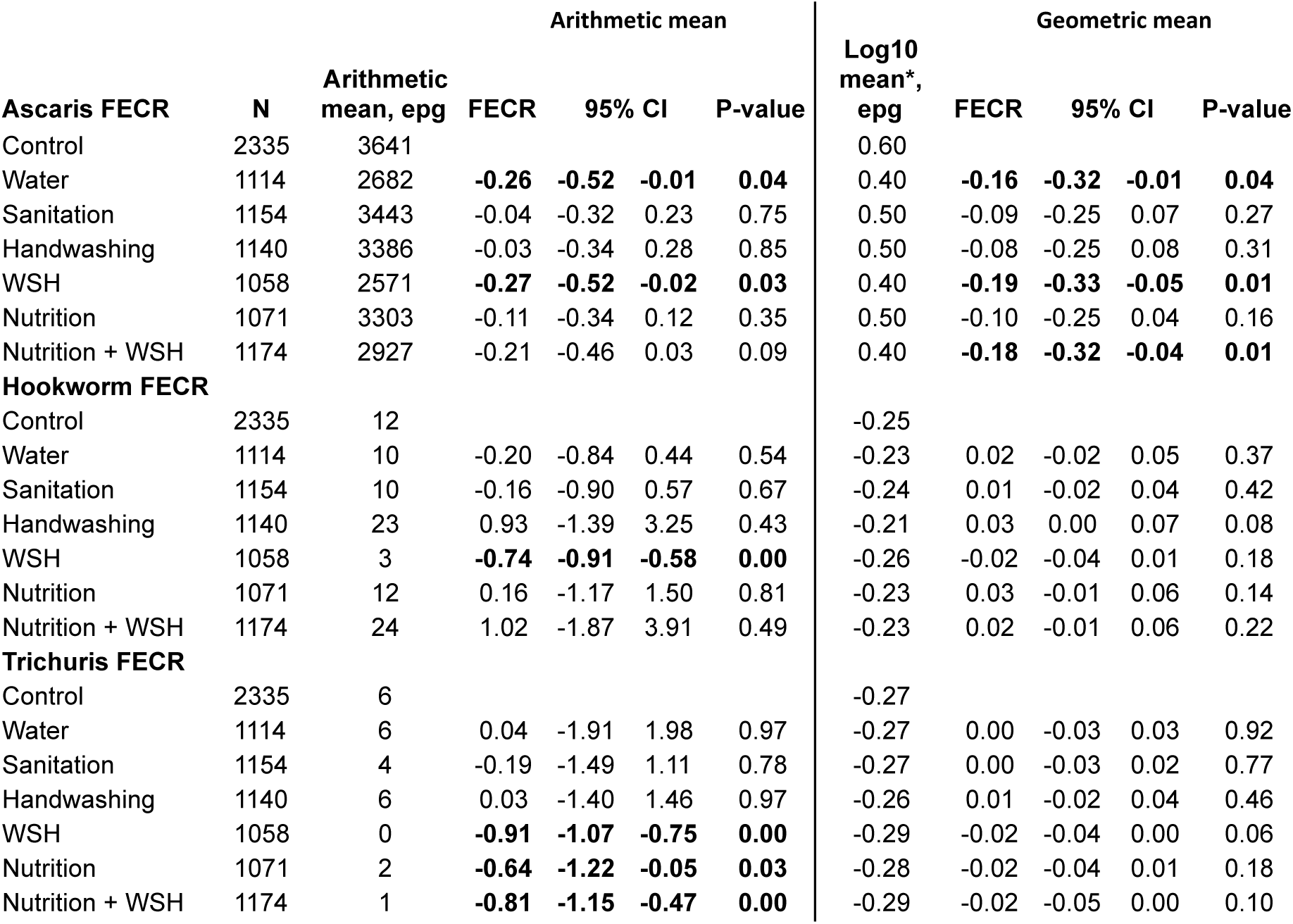
Effect of the interventions on infection intensity, measured by fecal egg count reduction (FECR) with arithmetic and geometric means in eggs per gram (epg). FECR estimated by targeted maximum likelihood estimation. Bold indicates p<0.05. ^∗^Values of 0.5 epg substituted for samples below the detection limit to calculate log-transformed mean

The FECR with arithmetic means indicated that children in the WSH arm had lower intensity infections with hookworm (3 eggs per gram [epg] *vs.* 11 epg in control) (Table 1). In addition, the FECR with arithmetic means indicated lower *Trichuris* infection intensity in the WSH (0 epg *vs.* 6 epg in control), nutrition (2 epg), and the WSHN (1 epg) arms. Children that received the WSHN intervention had 27% lower prevalence of coinfection with STH and *Giardia* compared to the control group (PR: 0.73, 95% CI 0.56, 0.97)(Table S4). STH coinfection was rare (<2% in control arm) and at similarly low levels in interventions arms (Table S4).

### Adjusted models and subgroup analyses

Adjusted effect estimates were similar to unadjusted effects (Table S4). Subgroup analyses of intervention effects stratified by child age, reported soil consumption (index children only), number of people living in the compound, deworming (*Ascaris* only), and time since defecation did not show any strong effect modification (Table S6).

## Discussion

This study provides new evidence on the effect of improved water, sanitation, handwashing in the household, and nutrition interventions, alone and in combination, on the prevalence of infection with STH and *Giardia*. Our findings demonstrate that an integrated water, sanitation, and handwashing intervention targeting the household environment in rural Kenya reduced *Ascaris* infection prevalence by 22%, while a water treatment intervention reduced *Ascaris* infection by 18%. Almost identical effect estimates generated by analyzing stool samples with microscopy and qPCR in a subset of arms leant additional credibility to the overall results (Figure 3). In addition, we found that improved nutrition did not enhance the effectiveness of the WSH intervention. *Trichuris* and hookworm prevalence were too low to precisely assess intervention impact in this setting, and *Giardia* was unaffected by the interventions. Although the integrated WSH intervention did not succeed in improving child growth or reducing symptomatic diarrhea in this trial(22), our findings confirm that WSH can effectively interrupt environmental helminth transmission.

A limited number of randomized controlled trials (RCTs) have previously analyzed the effect of WSH interventions on STH infection. Two RCTs in rural India found no impact of community sanitation interventions on helminth infections; however, both studies reported low usage rates of toilets among intervention households(28,29). Several school-based RCTs combining deworming with handwashing promotion have reported significant reductions in *Ascaris* reinfection prevalence in China, Ethiopia, and Peru(17,30). A school-based integrated WSH intervention combined with deworming in rural Kenya also reduced the odds of *Ascaris* reinfection(31). While previous RCTs demonstrate the success of school-based deworming combined with hygiene promotion, our results contribute new evidence from a large, cluster-randomized trial that improving WSH in the household environment can reduce *Ascaris* infections in a rural, low-income setting.

We did not detect an effect of the single sanitation intervention on STH infection prevalence. One potential explanation for the lack of impact may be that transitioning households from using traditional pit latrines to pit latrines with slabs may not have a measurable impact on STH transmission. A shift from households practicing open defecation to using latrines might be more likely to reduce STH transmission, with little additional benefit from improving latrine quality. A recent trial in Cote d’Ivoire reported greater reduction in hookworm infection prevalence among communities that received a community-led total sanitation intervention (designed to reduce open defecation levels) integrated with community-wide MDA compared to community-wide MDA alone (32). A second explanation may be that sanitation interventions are more effective at interrupting environmental transmission of pathogens when they are implemented at the community level(33), whereas our intervention only improved sanitation access in compounds with enrolled pregnant women.

The reductions in *Ascaris* prevalence in the combined arms could have resulted from improved water quality alone; *Ascaris* prevalence was 18% lower in the single water intervention arm than the control, a similar magnitude to the 22% reduction in the integrated intervention arms. Near identical reductions in *Ascaris* infection across all three water intervention arms suggests that water could have been an important transmission pathway in this population, which was interrupted by chlorine treatment. However, we cannot rule out contribution to reductions from other interventions in the combined arms; *Ascaris* prevalence was lower (20%) in each of the single sanitation, handwashing, and nutrition intervention arms, compared to 23% prevalence in the control arm. Chlorine is not known to inactivate *Ascaris* eggs, but one experimental study did find that chlorine can delay egg development and infectivity(34); it’s possible that delayed egg infectivity could reduce the risk of consuming an infective egg through drinking water. The proportion of households using jerry cans (a plastic water container with a narrow capped opening) to safely store drinking water was slightly higher in the water intervention arms than other arms (Tables S2 & S3). Our findings indicate that water is an understudied transmission pathway for *Ascaris*. We believe drinking water treatment should be further investigated as an STH control strategy, and that chlorine should be further explored as a method for inhibiting *Ascaris* egg development in drinking water supplies.

The combined WSHN intervention was similarly effective to WSH in reducing *Ascaris* prevalence, and improved nutrition did not reduce STH or *Giardia* infection on its own. Together, these results suggest that improved nutrition intervention did not reduce parasite infection in this population. Trials investigating the impact of micronutrient supplementation on STH infection or reinfection have reported mixed results(18). Our results are consistent with a Kenyan trial that found no effect of school-based micronutrient supplementation on reinfection with *Ascaris*(35). Considering interventions in this trial did not include treatment with antiparasitic drugs, further research would be valuable to understand if LNS supplementation could prevent parasite infections after drug treatment.

*Giardia* prevalence was unaffected by any of the interventions in this trial. Our results stand in contrast to results from the parallel WASH Benefits trial conducted in Bangladesh(36), which detected reductions in *Giardia* infection prevalence in the handwashing, sanitation, combined WSH, and combined WSHN arms(37). One potential explanation for lack of intervention effects in this trial is that water could be the primary transmission pathway for *Giardia* in this study setting, and *Giardia* is highly resistant to chlorination. The majority of households in the WASH Benefits Bangladesh trial accessed protected tubewells providing water with lower levels of fecal contamination compared to the springs and shallow wells accessed by households in this trial(27,38). Another potential explanation is that handwashing rates with soap were not high enough at the time of measurement to interrupt *Giardia* transmission; presence of soap and water at a handwashing station decreased from 78% at year one to 19% at year two among households in the WSH arm (Tables S2 & S3). *Giardia* is also zoonotic(4); exposure to avian and ruminant fecal contamination in the household environment could mitigate the effect of improved sanitation on transmission. Animal feces management was not a targeted behavior of the intervention packages.

This trial had some limitations. Chlorination does not inactivate protozoa, but was selected as the most appropriate water treatment intervention for the study context considering previous local acceptability, affordability, and effectiveness against bacterial and viral enteric pathogens. We measured parasite infections two years after intervention delivery; measurement among the study population at one year could have produced different results because of higher intervention adherence at that time (Table S2) and different child age-related exposures (e.g. younger children may be more likely to consume soil). We were unable to blind study participants due to the nature of the interventions; however, our outcomes were objective indicators of infection analyzed by blinded laboratory technicians, and blinded analysts replicated the data analysis.

During our trial, Kenya implemented a national school-based mass drug administration (MDA) program to reduce STH prevalence(39); and 43% of study children reported consuming deworming medication in the past 6 months (Table S6). Reported consumption of deworming medicine was similar across study arms, suggesting no systematic differences in program coverage or intensity between arms (Table S10). We observed similar *Ascaris* prevalence among study index children (23%, median age 2 years) and older children (22%, median age 5 years), suggesting that school-based MDA could be missing a key reservoir of infection among young, preschool aged children. Moreover, an environmental survey conducted during the national deworming program in our study region reported common detection of STH eggs in soil collected from the entrance to homes, with *Ascaris* eggs detected in soil in 19% of households(40). Taken together, these findings suggest additional control strategies beyond school-based deworming might be necessary to fully interrupt environmental STH transmission.

In contrast to most previous trials evaluating the effect of WSH or nutrition on STH infection, administering deworming medication was not included with our intervention. Our findings represent the potential impact of WSH and nutrition interventions in the context of exposure to a deworming program implemented at national scale. Although the magnitude of *Ascaris* prevalence reduction observed in the WSH and water intervention arms may be lower than what could be achieved by drug treatment in the short term, reduced STH infection after two years of intervention exposure indicates sustained impact. Our results support the proposal that improved WSH complement chemotherapy in the global effort to eliminate STH transmission.

## Acknowledgements

This research was financially supported in part by Global Development grant OPPGD759 from the Bill & Melinda Gates Foundation to the University of California, Berkeley. This manuscript was also made possible by the generous support of the American people through the United States Agency for International Development (USAID) (grant AID-OAA-F-13-00040 to Innovations for Poverty Action). The contents are the responsibility of the authors and do not necessarily reflect the views of USAID or the United States Government. We thank Charles Arnold for data management assistance, Michael Kremer for input on the study design, Araka Sylvie Biyaki for contributing to sample analysis, the WASH Benefits staff at IPA, and our study participants.

## References

1. Pullan RL, Smith JL, Jasrasaria R, Brooker SJ. Global numbers of infection and disease burden of soil transmitted helminth infections in 2010. Parasites & Vectors. Parasites & Vectors; 2014 Jan 21;7(1):1–19.

2. Rogawski ET, Bartelt LA, Platts-Mills JA, Seidman JC, Samie A, Havt A, et al. Determinants and Impact of Giardia Infection in the First 2 Years of Life in the MAL-ED Birth Cohort. J Pediatric Infect Dis Soc. Oxford University Press; 2017 Jun 1;6(2):153–60.

3. Bethony J, Brooker S, Albonico M, Geiger SM, Loukas A, Diemert D, et al. Soil-transmitted helminth infections: ascariasis, trichuriasis, and hookworm. The Lancet. 2006 May;367(9521):1521–32.

4. Certad G, Viscogliosi E, Chabé M, Cacciò SM. Pathogenic Mechanisms of Cryptosporidium and Giardia. Trends in Parasitology. Elsevier Ltd; 2017 Mar 20;:1–16.

5. Dickson R, Awasthi S, Williamson P, Demellweek C, Garner P. Effects of treatment for intestinal helminth infection on growth and cognitive performance in children: systematic review of randomised trials. BMJ. 2000 Jun 24;320(7251):1697–701.

6. Taylor-Robinson DC, Maayan N. Deworming drugs for soil-transmitted intestinal worms in children: effects on nutritional indicators, haemoglobin and school performance. … Database Syst Rev. 2012.

7. Donowitz JR, Alam M, Kabir M, Ma JZ, Nazib F, Platts-Mills JA, et al. A Prospective Longitudinal Cohort to Investigate the Effects of Early Life Giardiasis on Growth and All Cause Diarrhea. Clinical Infectious Diseases. 2016 Aug 24;63(6):792–7.

8. Simsek Z, Zeyrek FY, Kurcer MA. Effect of Giardia infection on growth and psychomotor development of children aged 0-5 years. J Trop Pediatr. 2004 Apr;50(2):90–3.

9. Prado MS, Cairncross S, Strina A, Barreto ML, Oliveira-Assis AM, Rego S. Asymptomatic giardiasis and growth in young children; a longitudinal study in Salvador, Brazil. Parasitology. Cambridge University Press; 2005 Jul 1;131(1):51–6.

10. Berkman DS, Lescano AG, Gilman RH, Lopez SL, Black MM. Effects of stunting, diarrhoeal disease, and parasitic infection during infancy on cognition in late childhood: a follow-up study. The Lancet. 2002;359(9306):564–71.

11. Owada K, Nielsen M, Lau CL, Clements ACA, Yakob L, Soares Magalhaes RJ. Measuring the Effect of Soil-Transmitted Helminth Infections on Cognitive Function in Children: Systematic Review and Critical Appraisal of Evidence. Advances in Parasitology. 2017 Jan 1.

12. Clarke NE, Clements ACA, Doi SA, Wang D, Campbell SJ, Gray D, et al. Differential effect of mass deworming and targeted deworming for soil-transmitted helminth control in children: a systematic review and meta-analysis. The Lancet. 2017 Jan;389(10066):287–97.

13. Jia T-W, Melville S, Utzinger J, King CH, Zhou X-N. Soil-Transmitted Helminth Reinfection after Drug Treatment: A Systematic Review and Meta-Analysis. Cooper PJ, editor. PLoS Negl Trop Dis. 2012 May 8;6(5):e1621.

14. Campbell SJ, Nery SV, McCarthy JS, Gray DJ, Magalhães RJS, Clements ACA. A Critical Appraisal of Control Strategies for Soil-Transmitted Helminths. Trends in Parasitology. Elsevier; 2016 Feb 1;32(2):97–107.

15. Savioli L, Smith H, Thompson A. Giardia and Cryptosporidium join the “Neglected Diseases Initiative.” Trends in Parasitology. 2006 May;22(5):203–8.

16. Speich B, Croll D, Fürst T, Utzinger J, Keiser J. Effect of sanitation and water treatment on intestinal protozoa infection: a systematic review and meta-analysis. The Lancet Infectious Diseases. Elsevier; 2016 Jan 1;16(1):87–99.

17. Strunz EC, Addiss DG, Stocks ME, Ogden S, Utzinger J, Freeman MC. Water, Sanitation, Hygiene, and Soil-Transmitted Helminth Infection: A Systematic Review and Meta-Analysis. Hales S, editor. PLoS medicine. Public Library of Science; 2014 Mar 25;11(3):e1001620–38.

18. Yap P, Utzinger J, Hattendorf J, Steinmann P. Influence of nutrition on infection and re-infection with soil-transmitted helminths: a systematic review. Parasites & Vectors. BioMed Central; 2014 May 19;7(1):229.

19. David LA, Maurice CF, Carmody RN, Gootenberg DB, Button JE, Wolfe BE, et al. Diet rapidly and reproducibly alters the human gut microbiome. Nature. Nature Publishing Group; 2015 Apr 9;505(7484):559–63.

20. Glendinning L, Nausch N, Free A, Taylor DW, Mutapi F. The microbiota and helminths: sharing the same niche in the human host. Parasitology. 2014 Sep;141(10):1255–71.

21. Arnold BF, Null C, Luby SP, Unicomb L, Stewart CP, Dewey KG, et al. Cluster-randomised controlled trials of individual and combined water, sanitation, hygiene and nutritional interventions in rural Bangladesh and Kenya: the WASH Benefits study design and rationale. BMJ Open. 2013 Aug 1;3(8):e003476–6.

22. Null C, Stewart CP, Pickering AJ, Global HDTL, 2018. Effects of water quality, sanitation, handwashing, and nutritional interventions on diarrhoea and child growth in rural Kenya: a cluster-randomised controlled trial. The Lancet Global Health. 2018.

23. Pilotte N, Papaiakovou M, Grant JR, Bierwert LA, Llewellyn S, McCarthy JS, et al. Improved PCR-Based Detection of Soil Transmitted Helminth Infections Using a Next-Generation Sequencing Approach to Assay Design. Albonico M, editor. PLoS Negl Trop Dis. Public Library of Science; 2016 Mar;10(3):e0004578.

24. Gruber S, van der Laan MJ. tmle: An R Package for Targeted Maximum Likelihood Estimation. Journal of Statistical Software. 2012 Nov 6;:1–35.

25. Balzer LB, van der Laan MJ, Petersen ML. Adaptive pre-specification in randomized trials with and without pair-matching. Statistics in Medicine. Wiley-Blackwell; 2016 Nov 10;35(25):4528–45.

26. Schulz KF, Grimes DA. Multiplicity in randomised trials I: endpoints and treatments. Lancet. Elsevier; 2005 May;365(9470):1591–5.

27. Pickering AJ, Arnold BF, Dentz HN, Colford JM, Null C. Climate and Health Co-Benefits in Low-Income Countries: A Case Study of Carbon Financed Water Filters in Kenya and a Call for Independent Monitoring. Environ Health Perspect. National Institute of Environmental Health Science; 2017 Mar;125(3):278–83.

28. Patil SR, Arnold BF, Salvatore AL, Briceno B, Ganguly S, Colford JM, et al. The Effect of India’s Total Sanitation Campaign on Defecation Behaviors and Child Health in Rural Madhya Pradesh: A Cluster Randomized Controlled Trial. Hunter PR, editor. PLoS medicine. Public Library of Science; 2014 Aug 26;11(8):e1001709.

29. Clasen T, Boisson S, Routray P, Torondel B, Bell M, Cumming O, et al. Effectiveness of a rural sanitation programme on diarrhoea, soil-transmitted helminth infection, and child malnutrition in Odisha, India: a cluster-randomised trial. The Lancet Global Health. Elsevier; 2014 Nov 1;2(11):e645–53.

30. Mahmud MA, Spigt M, Bezabih AM, Pavon IL, Dinant G-J, Velasco RB. Efficacy of Handwashing with Soap and Nail Clipping on Intestinal Parasitic Infections in School-Aged Children: A Factorial Cluster Randomized Controlled Trial. Bhutta ZA, editor. PLoS medicine. 2015 Jun 9;12(6):e1001837–16.

31. Freeman MC, Clasen T, Brooker SJ, Akoko DO, Rheingans R. The Impact of a School-Based Hygiene, Water Quality and Sanitation Intervention on Soil-Transmitted Helminth Reinfection: A Cluster-Randomized Trial. American Journal of Tropical Medicine and Hygiene. The American Society of Tropical Medicine and Hygiene; 2013 Nov 6;89(5):875–83.

32. Hürlimann E, Silué KD, Zouzou F, Ouattara M, Schmidlin T, Yapi RB, et al. Effect of an integrated intervention package of preventive chemotherapy, community-led total sanitation and health education on the prevalence of helminth and intestinal protozoa infections in Côte d’Ivoire. Parasites & Vectors. BioMed Central; 2018 Feb 27;11(1):115.

33. Harris M, Alzua ML, Osbert N, Pickering A. Community-Level Sanitation Coverage More Strongly Associated with Child Growth and Household Drinking Water Quality than Access to a Private Toilet in Rural Mali. Environmental Science & …. American Chemical Society; 2017 Jun 1;51(12):7219–27.

34. Oh K-S, Kim G-T, Ahn K-S, Shin S-S. Effects of Disinfectants on Larval Development of Ascaris suum Eggs. Korean J Parasitol. 2016 Feb;54(1):103–7.

35. Olsen A, Thiong’o FW, Ouma JH, Mwaniki D, Magnussen P, Fleischer Michaelsen K, et al. Effects of multimicronutrient supplementation on helminth reinfection: a randomized, controlled trial in Kenyan schoolchildren. Transactions of the Royal Society of Tropical Medicine and Hygiene. 2003 Jan;97(1):109–14.

36. Luby SP, Rahman M, Arnold BF, Unicomb L, Ashraf S, Winch PJ, et al. Effects of water quality, sanitation, handwashing and nutritional interventions on diarrhoea and child growth in rural Bangladesh: A cluster randomized trial. The Lancet Global Health.

37. Lin A, Ercumen A, Benjamin-Chung J, Arnold BF, Das S, Haque R, et al. Effects of water, sanitation, handwashing, and nutritional interventions on child enteric protozoan infections in rural Bangladesh: A cluster-randomized controlled trial. Clin Infect Dis. 3rd ed. 2018 Apr 13;16:87.

38. Pickering AJ, Ercumen A, Arnold BF, Kwong LH, Parvez SM, Alam M, et al. Fecal Indicator Bacteria along Multiple Environmental Transmission Pathways (Water, Hands, Food, Soil, Flies) and Subsequent Child Diarrhea in Rural Bangladesh. Environmental Science & Technology. American Chemical Society; 2018 Jun 14;52(14):7928–36.

39. Okoyo C, Nikolay B, Kihara J, Simiyu E, Garn JV, Freeman MC, et al. Monitoring the impact of a national school based deworming programme on soil-transmitted helminths in Kenya: the first three years, 2012 – 2014. Parasites & Vectors. BioMed Central; 2016 Jul 25;9(1):408.

40. Steinbaum L, Njenga SM, Kihara J, Boehm AB, Davis J, Null C, et al. Soil-Transmitted Helminth Eggs Are Present in Soil at Multiple Locations within Households in Rural Kenya. Aroian RV, editor. PLoS ONE. Public Library of Science; 2016;11(6):e0157780.

